# Gut Enterochromaffin Cells are Critical Drivers of Visceral Pain and Anxiety

**DOI:** 10.1101/2022.04.04.486775

**Authors:** James R. Bayrer, Joel Castro, Archana Venkataraman, Kouki K. Touhara, Nathan D. Rossen, Ryan D. Morrie, Aenea Hendry, Jessica Madden, Kristina N. Braverman, Gudrun Schober, Mariana Brizuela, Carla Bueno Silva, Holly A. Ingraham, Stuart M. Brierley, David Julius

## Abstract

Gastrointestinal (GI) discomfort is a hallmark of most gut disorders and represents a significant component of chronic visceral pain ^1^. For the growing population afflicted by irritable bowel syndrome (IBS), GI hypersensitivity and pain persist long after signs of tissue injury have resolved ^2^. IBS also exhibits a strong sex bias afflicting women three-fold more than men ^1^. Identifying the molecules, cells, and circuits that mediate both the acute and persistent phases of visceral pain is a critical first step in understanding how environmental and endogenous factors produce long-term changes in the nervous system or associated tissues to engender chronic pain syndromes ^3,4^. Enterochromaffin (EC) cells within the gut epithelium are exceedingly rare sensory neuroendocrine cells that detect and transduce noxious stimuli to nearby nerve endings via serotonin. Here, we manipulate murine EC cell activity using genetic strategies to ascertain their contributions to visceral pain. We show that acute EC cell activation is sufficient to elicit hypersensitivity to gut distension and necessary for the sensitizing actions of isovalerate, a bacterially derived short-chain fatty acid irritant associated with inflammatory GI disorders. Remarkably, prolonged EC cell activation by itself is sufficient to produce persistent visceral hypersensitivity, even in the absence of an instigating inflammatory episode. Perturbing the activity of these rare EC cells led to a marked increase in anxiety-like behaviors that normalized after blocking serotonergic signaling. Sex differences were also observed accross a range of assays indicating that females have a higher baseline visceral sensitivity. Our findings validate a critical role for EC cell-mucosal afferent signaling in acute and persistent GI pain while highlighting mechanistically defined genetic models for studying visceral hypersensitivity, sex differences, and associated behaviors.

## MAIN TEXT

Sensory signals from the gut are transmitted to the central nervous system by primary afferent nerve fibers, including those that innervate the intestinal mucosa, where they interact with epithelial cells that form the barrier between the luminal compartment and lining surface of the GI tract ^3^. This thin epithelial sheet contains rare but functionally diverse enteroendocrine cells that detect various physiologic signals and release hormones and neurotransmitters to regulate nutrient absorption and digestion, motility, feeding behaviors, and sensory perception ^5–7^.

Enterochromaffin (EC) cells are a unique subtype of sensory enteroendocrine cells that detect environmental and endogenous stimuli capable of eliciting or exacerbating pain, including dietary irritants, microbial metabolites, inflammatory agents, mechanical distension, and stress-associated hormones and neurotransmitters ^5,8–12^. Once activated, these excitable EC cells release serotonin onto nearby 5-HT_3_ receptor-positive sensory nerve endings that transmit nociceptive signals to the spinal cord ^8^. However, in addition to EC cells, sensory nerve endings themselves (and possibly other enteroendocrine cell types) are capable of detecting noxious stimuli ^4,13^. Indeed, distinct spinal afferents innervate the gut, including those that respond to distension with low or high thresholds or over a wide dynamic range, as well as stretch-insensitive mucosal fibers. Thus, it remains to be established whether and to what extent EC cells and their neighboring mucosal afferents contribute to acute and/or persistent visceral pain.

Here, we address these issues by silencing or ectopically activating EC cells in mice and assessing consequences regarding colonic sensory nerve fiber activity, neuroanatomy, and behavior. Our results clearly demonstrate that activation of EC cells is both necessary and sufficient to enhance nerve fiber excitability with attendant increases in acute and chronic visceral hypersensitivity and pain. Importantly, we also show that EC cell perturbation is associated with increased anxiety-like behaviors, as commonly experienced by individuals with gastrointestinal hypersensitivity or sensory integration disorders affecting the gut-brain axis ^1^. These findings establish EC cells and neighboring mucosal afferents as critical initiators of persistent pain associated with IBS or other inflammatory or post-inflammatory GI disorders.

### EC cells transduce bacterial metabolite-evoked visceral hypersensitivity

Bacterially derived short-chain fatty acids are known activators of enteroendocrine cells ^8,14^. Some fatty acid species, such as isovalerate, are elevated in individuals with visceral hypersensitivity and may contribute to enhanced pain sensation ^14,15,16^. We have previously shown that isovalerate activates EC cells via the G protein-coupled receptor Olfr558, thereby producing serotonin-dependent mechanical hypersensitivity of primary afferents in an ex vivo nerve-gut model ^8^. Because both EC cells and sensory nerve fibers are mechanically responsive, we wondered whether isovalerate enhances sensitivity of nerve fibers to other, non-mechanical stimuli. Indeed, in ex vivo nerve-gut preparations from mice expressing channelrhodopsin in Na_V_1.8-positive primary afferents (Na_V_1.8-Ai32 mice), application of isovalerate boosted sensitivity of mucosal nerve fibers to light (Fig. 1a-c and Extended Data Fig1a-c). Thus isovalerate-mediated sensitization is not specific to mechanical stimulation of either EC cells or sensory neurons, but more broadly enhances excitability of primary afferent nerve fibers.

**Figure 1 -.**
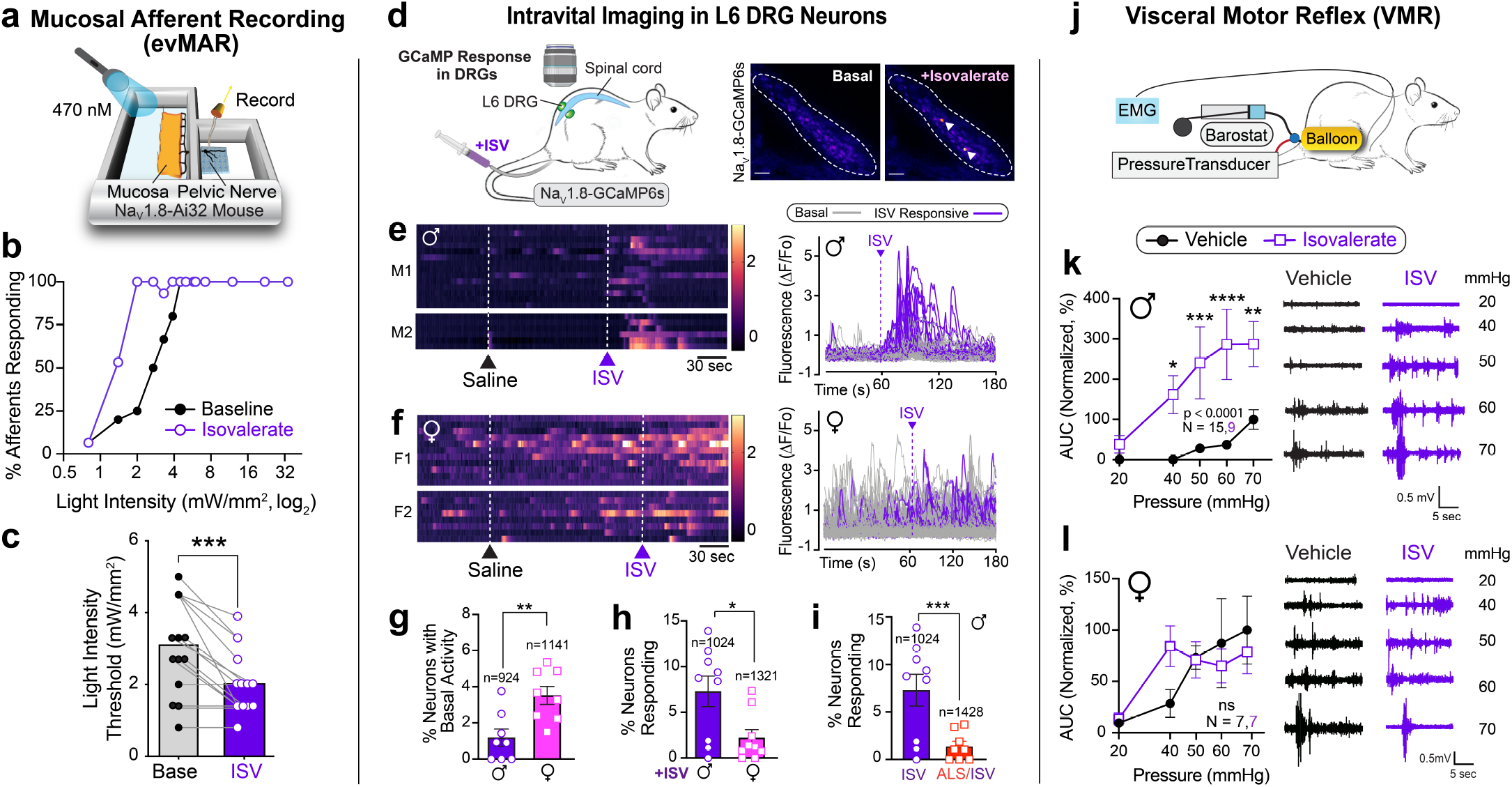
EC cells mediate sex-dependent response to irritants via serotonergic signaling. **a,** Illustration of the set-up used for the optogenetic ex vivo ‘flat sheet’ colon pelvic nerve mucosal afferent recordings from Na_V_1.8-Ai32 mice when stimulated with light (l=470 nm, 10 ms pulses at 1Hz with increasing intensity from 0.8 – 34 mW/mm^2^) plus vehicle or iso-valerate (200 μM). **b-c,** Percentage of neurons responding at indicated light intensities and activation thresholds of mucosal afferents before and after isovalerate treatment (n=15). **d,** Schematic of intravital calcium imaging and representative images of DRGs (outlined by dashed lines) at baseline and after intracolonic delivery of isovalerate; white arrowheads point to responsive neurons. **e-f,** Heatmaps from two representative males (e) or females (f) showing in vivo Na_V_1.8-GCaMP6s responses of L6 DRG neurons to intracolonic application of saline followed by an isovalerate bolus (200 μM) with examples of fluorescence changes shown in righthand panels (purple traces depict isovalerate sensitive neurons). **g,** Group data comparing percentage of neurons with basal activity in males versus females. **h,** Group data comparing percentage of DRG neurons responding in males versus females. **i,** Group data showing percentage of DRG neurons that respond to isovalerate (replotted from panel h) or isovalerate plus alosetron (10 μM), n = total neurons analyzed per condition. **j,** Schematic of experimental setup for measuring electromyography visceromotor response (VMR) to colorectal distension following intra-colonic administration of vehicle or isovalerate (200 μM). **k-l,** VMR responses for post-pubertal wild type healthy males (k) or females (l). Biological replicates (N’s) shown in graphs, and representative traces shown in respective righthand panels. Chi-square test (2.0 mW/ mm^2^: p < 0.001, 2.7 mW/mm^2^: p < 0.01) in panel b. Student t-test (paired, 2 tailed) in panel c. Student t-test (unpaired, 2 tailed) in panel g. Two-way ANOVA (Šidák’s multiple-comparisons test) in panels h and i. *p < 0.05, **p < 0.01, ***p < 0.001, ***p < 0.0001.

Intravital calcium imaging of dorsal root ganglia in Na_V_1.8-GCaMP6s mice allowed us to assess how many neurons are sensitized by isovalerate in vivo (Fig 1d). In males, nearly 7% of neurons within lumbosacral ganglia were activated by intracolonic administration of isovalerate (Fig. 1e). Remarkably, we noticed that basal neuronal activity was substantially higher in females, thereby diminishing the percentage of cells showing enhanced sensitivity following administration of isovalerate (Fig. 1f-h). In males, the isovalerate response was blocked by co-administration of the selective 5-HT_3_ receptor antagonist, alosetron (Fig. 1i and Extended Data Fig. 1d), consistent with a mechanism in which isovalerate elicits release of serotonin from EC cells, followed by activation of 5-HT_3_ receptors on nearby sensory nerve endings.

To determine whether isovalerate enhances behavioral pain responses, we performed a series of colorectal distensions in mice and measured reflex motor responses of abdominal muscles in the presence or absence of intracolonically-delivered isovalerate (Fig 1j). Indeed, isovalerate produced substantial enhancement of this visceromotor response (VMR) across a range of distension pressures in males (Fig. 1k and Extended Data Fig. 1e). This behavioral effect was severely blunted in females perhaps reflecting the narrower dynamic range (Fig 1l), in keeping with the higher cellular baseline activity noted above. Importantly, colonic compliance was the same irrespective of sex or treatment (Extended Data Fig. 1f).

We next asked if the sensitizing effect of isovalerate was mediated specifically by EC cells. To address this question, we exploited the ability of tetanus toxin to inhibit synaptic vesicle fusion and neurotransmitter release from excitable cells. We employed an intersectional genetic approach to target expression of tetanus toxin (or DREADD receptors, see below) to EC cells. Briefly, a *Pet1Flp* allele that favors recombination in gut enteroendocrine cells was used in conjunction with a Cre driver (*Tac1Cre*) that limits recombination to tachykinin-expressing cells (Fig. 2a). Indeed, within the gut our *Pet1Flp;Tac1Cre* line selectively targets EC cells (Fig. 2b and Extended Data Figs. 2a, 3a, 4a) and histological analyses showed no evidence of transgenic marker expression within serotonergic raphe nuclei, the spinal cord, or DRG sensory ganglia, consistent with gut-specific EC cell expression (Extended Data Fig. 3b-d). In addition to demonstrating selective targeting, these data highlight the remarkable rarity of EC cells and the relatively small number of intestinal cells that are targeted and controlled by this genetic manipulation. We were also able to target EC cells by using a *Vil1Cre* driver that limits recombination to the intestinal epithelium (Fig. 2b), and this mouse line was used in one set of ex vivo studies (see below).

**Figure 2 -.**
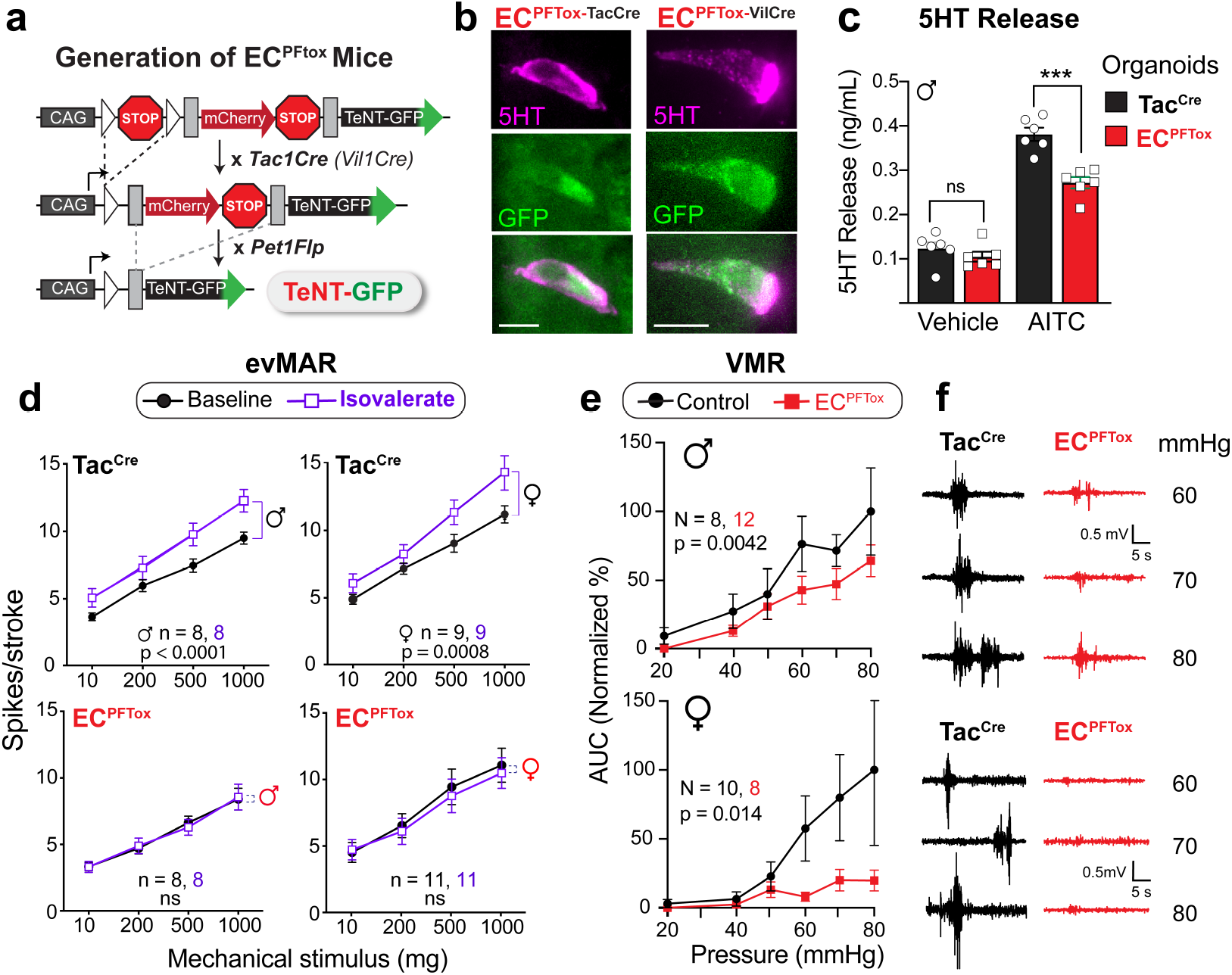
Silencing EC cells attenuates colonic sensitivity to irritants and pressure. **a, I**ntersectional genetic strategy to selectively express tetanus toxin (TeNT) in EC cells. **b,** Representative EC cells showing overlapping expression of TeNT (GFP) and 5-HT (magenta). Scale bars = 10 μm. **c,** 5-HT release as detected by ELISA is blunted in EC^PFTox^ intestinal organoids compared to Tac^Cre^ control organoids upon stimulation with allyl isothiocyanate (AITC, 20 μM; N = 6 per group). **d,** Group data from mucosal afferent recordings (evMAR) at baseline and following isovalerate (200 μM) application for Tac^Cre^ control and EC^PFTox^ male and female cohorts. **e,** VMR to colorectal distension is shown for male and female EC^PFTox^ mice compared to littermate controls. **f,** Representative traces for control Tac^Cre^ (left) or EC^PFTox^ (right) in male (upper) and female (lower) mice. Student’s t-test (unpaired, 2-tailed) in panel c, two-way ANOVA (Šidák’s multiple-comparisons test) in panels d and e, *** p < 0.001.

To validate a functional effect of tetanus toxin expression, we first measured stimulus-evoked serotonin release using intestinal organoids derived from the tetanus toxin expressing animals (EC^PFTtox^). The electrophilic irritant, allyl isothiocyanate (AITC), was used to activate native TRPA1 channels on EC cells and stimulate serotonin release. Indeed, we observed a substantial (~30%) diminution in AITC-evoked serotonin release from EC^PFTox^ compared to Tac^Cre^ control organoids (Fig. 2c). Moreover, whereas application of isovalerate to ex vivo gut-nerve preparations from Tac^Cre^ control animals enhanced responses of sensory nerves to mechanical stimulation of the mucosa (Fig. 2d and Extended Data Fig. 2b-d), no such sensitization was observed with preparations from male or female EC^PFTox^ mice (Fig. 2d and Extended Data Fig. 2b-d), suggesting that both EC cells and mucosal afferents are critical to generating visceral pain. In further support of this notion, we found that VMRs to colorectal distension were diminished in EC^PFTox^ mice versus control littermates (Fig. 2e,f and Extended Data Fig. 2e,f), especially in females, where the response was completely abrogated. Taken together, our results thus far show that EC cells contribute substantially to visceral nociception. Moreover, silencing EC cells has a proportionally greater effect in females. This, and the greater sensitivity of males to isovalerate, suggests that EC cells from females have a higher baseline activity and a narrower dynamic range, factors that may contribute to the sex bias in visceral sensitivity.

### Selective EC cell activation drives visceral hypersensitivity in a sex-dependent manner

Having substantiated a role for EC cells in visceral pain, we next used chemogenetics to ask if their activation is sufficient to generate acute and/or persistent visceral hypersensitivity. Using the same transgenic strategy described above, an activating DREADD receptor (hM3Dq) was expressed in EC cells to enable their selective pharmacological activation (EC^hM3Dq^, Fig. 3a,b and Extended Data Fig. 4a). Indeed, the application of a DREADD agonist, deschloroclozapine (DCZ) to intestinal organoids derived from these EC^hM3Dq^ mice promoted EC cell activation and serotonin release (Fig. 3c and Extended Data Fig. 4b). Furthermore, DREADD receptor activation rendered fibers from EC^hM3Dq^ animals hypersensitive to mucosal mechanical stimulation (Fig. 3d) without affecting those from Tac^Cre^ controls (Extended Data Fig. 4c-e). Here again, sensitization was more pronounced in males, consistent with the idea that basal EC cell activity is elevated in females compared to males, thereby diminishing the dynamic range of their response to ectopic stimuli.

**Figure 3 -.**
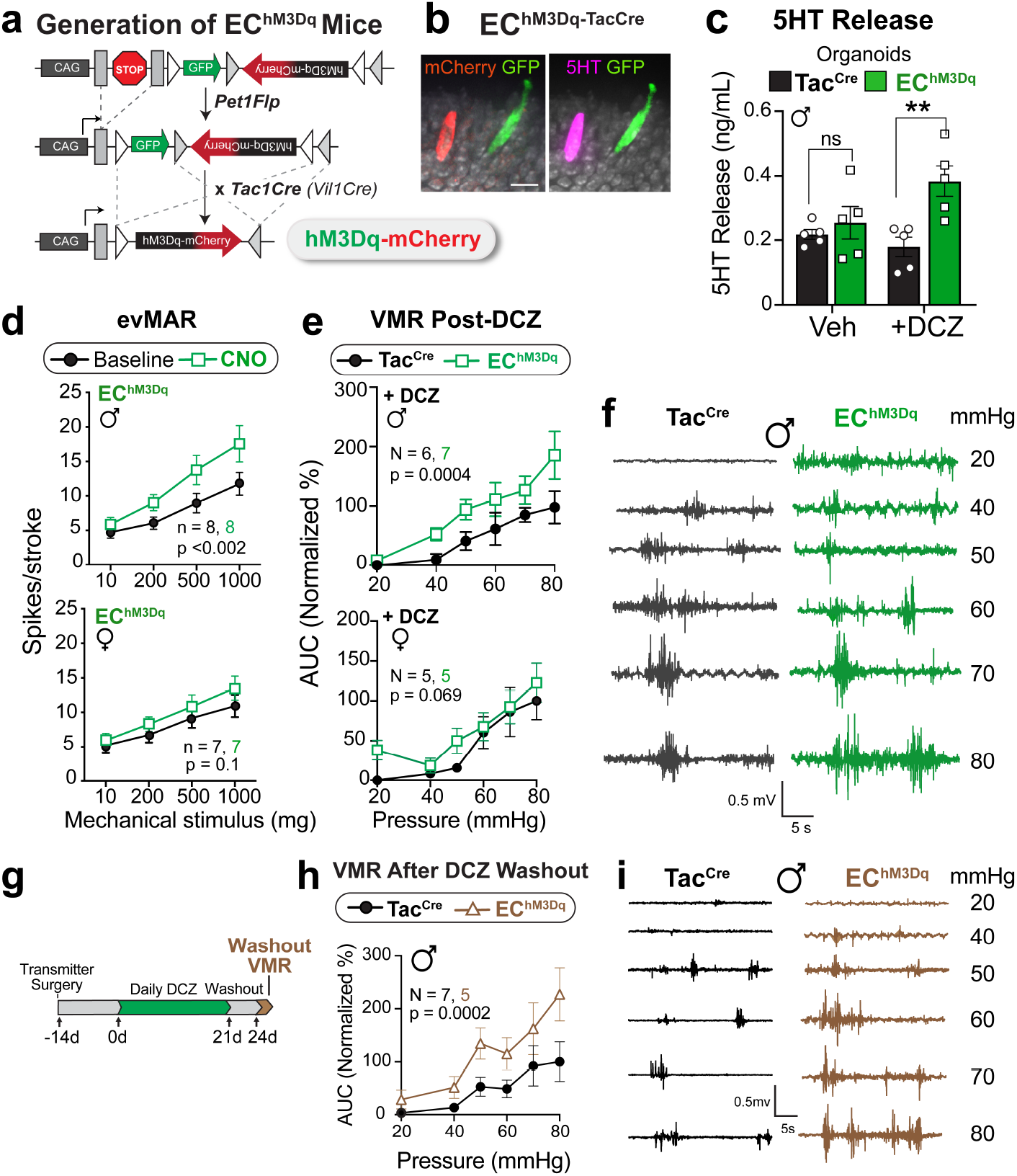
Activating EC cells increases afferent output in males and induces long-term hypersensitivity. **a,** Intersectional genetic strategy for expressing DREADD-hM3Dq in EC cells. **b,** Representative EC cell showing DREADD (mCherry) and 5-HT (magenta) co-expression. Scale bar = 15 μm. **c,** 5-HT release from Tac^Cre^ control or EC^hM3Dq^ intestinal organoids following treatment with deschloroclozapine (DCZ, 1.7 μM). **d,** Group data from mucosal afferent recordings (evMAR) for EC^hM3Dq^ mice at baseline and following clozapine N-oxide (CNO, 100 μM) application showing sensitization in males, but not females. **e,** VMR data following acute stimulation with DCZ (75 μg/kg i.p., 15 min) in male (upper) and female (lower) Tac^Cre^ and EC^hM3Dq^ cohorts. **f,** Representative EMG traces from a Tac^Cre^ control (left) and EC^hM-3Dq^ (right) male post-DCZ treatment. **g,** Timeline for chronic administration of DCZ (75 μg/kg, i.p.) for 21 days followed by a 3-day washout. **h,** VMR data showing persistent visceral hypersensitivity in male EC^hM3Dq^ mice compared to Tac^Cre^ controls following DCZ washout. **i,** Representative traces at each distension pressure for a male Tac^Cre^ and EC^hM3Dq^ mouse. Student’s t-test (unpaired, 2-tailed) in panel c; two-way ANOVA (Šidák’s multiple-comparisons test) in panels d, e, h. **p < 0.01.

**Figure 4 -.**
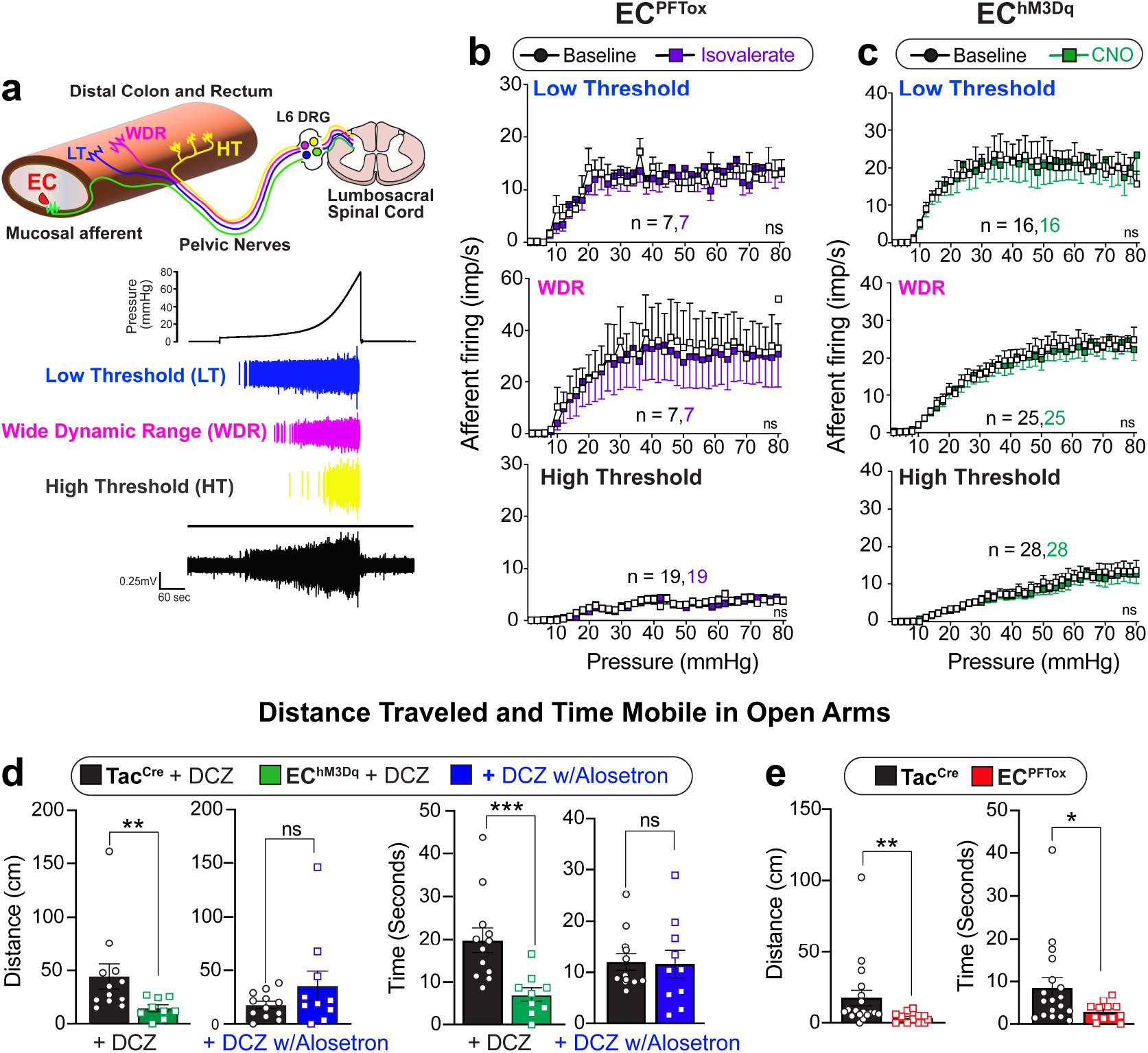
Manipulating EC cell-mucosal afferent activity induces anxiety-like behavior. **a,** Graphical illustration of different afferent subtypes innervating the colon and representative ex vivo ‘intact’ colonic afferent recordings showing low threshold (LT), wide dynamic range (WDR), and high threshold (HT) distension sensitive afferents from the pelvic nerve. **b-c,** Group data showing afferent firing to increasing distension pressures in preparations from EC^PFTox^ mice (N = 3 female and 2 male) at baseline and following isovalerate or EC^hM3Dq^ mice (N = 7 female and 2 male) at baseline and following CNO, as indicated. **d-e,** Distance traveled and time mobile in open arms of an elevated plus-maze using naive male and female cohorts (combined) comparing Tac^Cre^ control mice to either EC^hM3Dq^ or EC^PFTox^ mice 15 minutes post-DCZ (75 μg/kg i.p.) with or without alosetron (10 μM), as indicated. Two-way ANOVA (Šidák’s multiple-comparisons test) for panels b and c. Student t-test (unpaired Mann Whitney U test, 2-tailed) for panels d and e. *p < 0.05, **p < 0.01, ***p < 0.001.

Does this sensitization of nerve fibers also translate to visceral hypersensitivity? To address this question, we administered a DREADD agonist to EC^hM3Dq^ mice 15 min prior to colorectal distension, in which case we observed a significant increase in the VMR across a range of colonic pressures for males (Fig. 3e,f and Extended Data Fig. 4f), with a trend towards significance in females. This sex difference is again consistent with the notion that EC cells in females have a higher tonic contribution to colonic sensitivity and thus a smaller window for enhancement.

### Prolonged EC activation results in persistent visceral hypersensitivity

Our results thus far show that stimulation of EC cells is sufficient to produce acute visceral hypersensitivity. But can prolonged EC cell activation mimic a chronic hypersensitive state resembling that observed in IBS? To address this question, we treated mice with a DREADD agonist daily for three weeks and assessed the VMR to colonic distension 72 hours after cessation of agonist treatment (Fig. 3g). Remarkably, EC^hM3Dq^ animals maintained heightened sensitivity compared with control animals following drug removal, demonstrating that EC cells are sufficient for driving both acute and persistent visceral pain (Fig. 3h, i).

Considering previous studies suggesting a role for EC cell-mediated serotonin release in regulating gut motility ^12,17^, we examined the relevant parameters in EC^hM3Dq^ and EC^PFTox^ mice, including whole GI and colonic transit times. No significant differences were observed in these tests or in serum serotonin levels (Extended Data Fig. 5) highlighting the specificity of the pain phenotypes observed with these transgenic models.

### Mucosal nerve fibers selectively transmit nociceptive signals from EC cells

Primary afferent nerve fibers terminate in distinct layers of the gut, including the mucosal, muscular, and vascular compartments, as well as the mesentery (Fig. 4a) ^18^. It has generally been assumed that nerve fibers with low and high thresholds to distension drive visceral pain, rather than distension-insensitive mucosal afferents, which are instead exquisitely sensitive to small deformations of the mucosa ^19^. However, given the observed necessity of EC cells in mediating this pain response, we asked whether fibers other than those innervating the mucosa are impacted by EC cell activation. To address this question, we used a modified ex vivo nerve-gut model in which pressurization of an intact colonic loop activates afferent spiking in distension-sensitive afferents (including low and high threshold and wide dynamic range fibers) that detect circumferential tension applied across their receptive fields. Nerves innervating the mucosal layer do not respond in this paradigm, allowing us to distinguish them from these other afferent types. These recordings were performed with mice in which recombination was directed by *Pet1Flp;Vil1Cre*.

In Vil^Cre^ control mice, application of either isovalerate or a DREADD agonist failed to enhance activation of a variety of distension-sensitive afferents that innervate the non-mucosal layers (Extended Data Fig. 6a). As expected, preparations from EC^PFTox^ female or male mice, which showed diminished VMRs to colorectal distension, did not affect distension-sensitive nerve fiber activation before or after treatment with isovalerate (Fig. 4b). The same was true for preparations from EC^hM3Dq^ female and male mice, where no effect of DREADD agonist treatment was observed for the population of distension-sensitive nerve fibers (Fig. 4c). Taken together, these results suggest that modulation of visceral pain can be driven by distension-insensitive mucosal afferents that are instead sensitive to local tissue deformation and interact directly with EC cells in the mucosal layer.

### EC cell-mucosal afferent signaling axis modulates anxiety

Functional GI disorders are often associated with increased anxiety as a manifestation of complex pain or sensory processing disorders that might affect bidirectional gut-brain signaling ^1^. We therefore asked whether chronic activation or suppression of EC cells influences anxiety states in these animals by assessing behavior in a standard elevated plus maze (EPM) paradigm. We found that naive EC^hM3Dq^ animals treated with DCZ spent less time in the open arms (although total movement parameters in the closed arms did not differ), indicative of increased anxiety (Fig. 4d and Extended Data Fig 6b). These behavioral responses were equivalent in males and females.

To determine if increased anxiety in DCZ-treated EC^hM3Dq^ mice results from EC cell-mediated serotonin release and consequent activation of 5-HT_3_ receptors on primary mucosal afferents, these assays were repeated with co-administration of the 5-HT_3_ receptor antagonist, alosetron. Alosetron reversed the effects of DCZ (Fig. 4d), supporting the importance of serotonergic EC cell signaling in modulating both nociceptive and affective components of disordered gut-brain communication. Interestingly, EC^PFTox^ animals also spent less time in the open arms, but not in the closed arms (Fig 4e and Extended Data Fig 6c), suggesting that perturbation of GI function through chronic suppression of EC cell activity is similarly anxiogenic. This was further substantiated by a significant change in the marble-burying assay in contrast to normal nestlet building or conditional fear behavior (Extended Data Fig. 6d-f).

## DISCUSSION

Recent studies suggest that mediators released from the colonic epithelium can elicit ^20,21^ or inhibit ^22,23^ visceral pain. Our data now show that visceral pain and anxiety can be driven by an exceedingly small subpopulation of enteroendocrine cells, the EC cells, and their interaction with mucosal afferents. This is remarkable as mucosal afferents, which respond only to the smallest of stimuli deforming the mucosa, have not previously been considered key drivers of visceral pain, even though they show heightened responses in models of chronic visceral hypersensitivity ^24^. This role has historically been considered the domain of low threshold ^25^, wide dynamic range ^25^ or high threshold ^3,4,26^ distension-sensitive afferents, which are polymodal and respond to a wide variety of inflammatory and immune mediators ^13^. Our findings add to the ever-increasing contribution of EC cells to a myriad of gut functions ^7,27^ such as mechano-^12,28,29^ and chemo-sensation ^5,6,8–11,30,31^. How can EC cells assume such functional diversity? The answer may reflect their location within specific segments of the gastrointestinal tract ^32,33^ and the neuronal subtypes with which they interact, including those of vagal ^10,30,31^, spinal ^8,9^, or enteric origin ^12^. This study also supports a crucial role for serotonin and 5-HT_3_ receptors at the interface between EC cells and mucosal afferents. Alosetron, which is approved for the treatment of severe IBS with diarrhea ^34^, has been proposed to exert its analgesic effect centrally ^35,36^ or through inhibition of high threshold nociceptors within the gut wall ^37^. Our results now suggest an alternative or additional mechanism to account for this analgesic effect.

Irritable bowel syndrome is often initiated by profound intestinal inflammation, but pain and hypersensitivity persist long after resolution of tissue damage ^1,2^, likely reflecting a multifactorial process involving disruption of mucosal integrity with effects on both the epithelium and underlying sensory nerve fibers. However, our results now suggest that chronic activation of EC cells is, by itself, sufficient to generate persistent visceral hypersensitivity, providing a targeted experimental approach to study this maladaptive response in the absence of other confounding pathophysiologic variables. Moreover, our findings suggest that EC cells represent a relevant therapeutic target for ameliorating chronic visceral pain.

Functional GI disorders are especially prevalent among women^1^. Our data suggest that EC cells contribute more substantially to visceral pain in female versus male mice, perhaps reflecting a greater tonic input of these cells to nociception in females. Understanding the sex-specific molecular and functional differences in EC cells may provide mechanistic insight into the well-recognized sex disparity in visceral pain syndromes. Functional GI disorders are also associated with increased anxiety, and treatment with selective serotonin reuptake inhibitors may have a dual benefit in targeting both peripheral and central sites ^38^. Interestingly, we found that both male and female mice display enhanced anxiety with EC cell inhibition or activation, suggesting that this integrative behavioral state is an especially sensitive readout of visceral dysregulation while further highlighting the importance of the gut-brain axis in neurogastroenterological disorders ^39^.

## METHODS

### Ethics

Experiments were approved and performed in accordance with the guidelines of the Animal Ethics Committees of the South Australian Health and Medical Research Institute (SAHMRI), the University of South Australia and Flinders University, and with UCSF Institutional Animal Care Committee guidelines, the National Institutes of Health Guide for Care and Use of Laboratory Animals, and recommendations of the International Association for the Study of Pain.

### Generation of transgenic mouse lines

Excitatory DREADD hM3Dq expression was directed to enteroendocrine cells using the Cre and Flp dependent hM3Dq reporter mouse, *RC::FL-hM3Dq* (Jackson Labs, Strain #026942). This mouse line was crossed with the transgenic *Pet1Flp* line ^40^ (gift of Dr. Susan Dymecki), which expresses Flp recombinase under the control of the ePet1 transcription factor. ePet1 is expressed in serotonergic neurons in the brain ^40^, and enteroendocrine cells in the intestines ^41^. Mice hemizygous for *Pet1Flp* and homozygous for *RC::FL-hM3Dq* were then crossed to homozygous *Tac1Cre* (Jackson Labs, strain #021877) mice to achieve EC cell-specific expression of the DREADD reporter in EC^hM3Dq^ mice. In addition to EC cells, a sparse population of neurons in the medullary raphe has been reported to be Tac1+/Pet+ ^42^. Therefore, to look at intestine-specific effects of the DREADD receptor activation in enteroendocrine cells, *Pet1Flp* hemizygous, *RC::FL-hM3Dq* homozygous mice were bred to *Vil1Cre* (Jackson Labs, Strain #021504) hemizygous, *RC::FL-hM3Dq* homozygous mice. *Vil1Cre* directs expression of Cre recombinase exclusively to the intestinal epithelium ^43^. To inhibit serotonin release from EC cells, we targeted expression of the tetanus toxin light chain to enteroendocrine cells using the Cre and Flp dependent tetanus toxin light chain reporter mouse, *RC::PFTox* ^44^ (gift of Dr. Susan Dymecki). The same breeding scheme was deployed as above, i.e., *Pet1Flp* hemizygous/*RC::PFTox* homozygous mice x *Tac1Cre* homozygous mice to generate EC^PFTox^ mice and *Pet1Flp* hemizygous/*RC::PFTox* homozygous mice x *Vil1Cre* hemizygous/*RC::PFTox* homozygous mice. Of note, the above breeding schemes were required to avoid germline recombination of Cre reporter alleles by *Tac1Cre*.

### In vivo assessment of pain-related behavior

We used abdominal EMG to monitor the visceromotor response (VMR) to colorectal distension (CRD) in fully awake animals ^22,26,45^. For C57/BL6 and EC^PFTox^ experiments, the bare endings of two Teflon-coated stainless steel wires (Advent Research Materials Ltd, Oxford, UK) were sutured into the right abdominal muscle and tunneled subcutaneously to be exteriorized at the base of the neck for future access ^22,26,45^. At the end of the surgery, mice received prophylactic antibiotics (C57/BL6 animals; Baytril; 5 mg/kg s.c.) and analgesic (buprenorphine; 0.4 mg/10 kg s.c.), then they were housed individually and allowed to recover for at least three days before assessment of VMR. For EC^hM3Dq^ experiments, a wireless transmitter was implanted to enable repeated measurements over time (ETA-F10; Data Sciences International, New Brighton, MN USA). The transmitter was placed subcutaneously, and leads were implanted into the abdominal musculature as above. Animals were allowed to recover for at least seven days before undergoing baseline VMR. On the day of VMR assessment, mice were briefly anesthetized using isoflurane before receiving a 100 μL enema of vehicle (sterile saline) or isovalerate (200 μM). A lubricated balloon (2 cm length) was gently introduced through the anus and inserted into the colorectum up to 0.25 cm past the anal verge. The balloon catheter was secured to the base of the tail and connected to a barostat (Isobar 3, G&J Electronics, Willowdale, Canada) for graded and pressure-controlled balloon distension. For EC^hM3Dq^ experiments, DCZ (75 μg/kg i.p.) was administered 15 min prior to first distension. Mice were allowed to recover from anesthesia in a restrainer with dorsal access for 15 min prior to initiation of the distension sequence. Distensions were applied at 20-40-50-60-70-80mmHg (20 s duration each) at 2 min intervals so that the last distension was performed ~20 min after intracolonic treatment with either vehicle or isovalerate and ~40 min of DCZ administration. Following the final distension, colonic compliance was assessed by applying graded volumes (40–200 μL, 2 s duration each) to the balloon in the colorectum of fully awake mice while recording the corresponding colorectal pressure as described previously ^22^,^26^,^45^. For the VMR recordings, the EMG electrodes were relayed to a data acquisition system. The signal was recorded (NL100AK headstage), amplified (NL104), filtered (NL 125/126, Neurolog, Digitimer Ltd, bandpass 50–5000 Hz), and digitized (CED 1401, Cambridge Electronic Design (CED), Cambridge, UK) or for EC^hM3Dq^ experiments through the Ponemah Software System (Data Sciences International) for off-line analysis using Spike2 Software (CED). The analog EMG signal was rectified and integrated. To quantify the magnitude of the VMR at each distension pressure, the area under the curve (AUC) during the distension (20 s) was corrected for baseline activity (AUC pre-distension, 20 s). We also calculated the total AUC, the summation of data points across all distension pressures for each animal.

### Colon-pelvic nerve preparation for flat-sheet single fiber mucosal afferent recordings

Male and female wild-type or transgenic EC^hM3Dq^ or EC^PFTox^ mice were humanely killed by CO2 inhalation. The colon and rectum with attached pelvic nerves were removed, and recordings from mucosa afferents were performed as previously described 8,19. Briefly, the colon was removed and pinned flat, mucosal side up, in a specialized organ bath. The colonic compartment was superfused with a modified Krebs solution (in mM: 117.9 NaCl, 4.7 KCl, 25 NaHCO3, 1.3 NaH2PO4, 1.2 MgSO4 (H2O)7, 2.5 CaCl2, 11.1 D-glucose), bubbled with carbogen (95% O2, 5% CO2) at a temperature of 34°C. All preparations contained the L-type calcium channel antagonist nifedipine (1 μM) to suppress smooth muscle activity and the prostaglandin synthesis inhibitor indomethacin (3 μM) to suppress potential inhibitory actions of endogenous prostaglandins. The pelvic nerve bundle was extended into a paraffin-filled recording compartment in which finely dissected strands were laid onto a mirror, and a single fiber placed on the platinum recording electrode. Action potentials generated by mechanical stimuli to the colon’s receptive field, pass through the fibers into a differential amplifier, filtered, sampled (20 kHz) using a 1401 interface (CED, Cambridge, UK), and stored on a PC for off-line analysis. Categorization of afferents properties followed our previously published classification system ^19^. Receptive fields were identified by systematically stroking the mucosal surface with a still brush to activate all subtypes of mechanoreceptors. In short, pelvic mucosal afferents respond to delicate mucosal stroking (10 mg von Frey hairs; vfh) but not circular stretch (5 g). Stimulus-response functions were then constructed by assessing the total number of action potentials generated in response to mucosal stroking with 10, 200, 500, and 1000 mg vfhs. For wild type and transgenic EC^PFTox^ mice, isovalerate (200 μM) was applied to the mucosal epithelium for 15 min via a small metal ring placed over the receptive field of interest and mechanosensitivity re-tested. For wild type and transgenic EC^hM3Dq^ mice, the DREADD agonist CNO (100 μM) was applied to the mucosal epithelium for 15 min via a small metal ring placed over the receptive field of interest and mechanosensitivity re-tested.

### Optogenetic stimulation of colonic mucosal afferents

As Scn10a (Na_V_1.8) is expressed by >95% of colonic afferents 46 but not by ECs 8, we crossed Scn10a-Cre mice (gift from Dr. Wendy Imlach, Monash University, Australia) with Ai32 (ChR2)-floxed mice (JAX strain #012569; purchased from The Jackson Laboratory; Bar Harbor, ME, USA) to produce Na_V_1.8-Ai32 mice. This allowed us to optogenetically activate colonic mucosal afferents in an EC-independent manner. Colonic recordings were prepared from Na_V_1.8-Ai32 mice using methods described above. Once mucosal afferents had been identified by their responsiveness to fine mucosal stroking (10 mg vfh), but not to circular stretch (5 g), afferents were left to rest for 10 minutes before optogenetic stimulation. Afferents were illuminated with a light wavelength of 470 nm in 10ms pulses at 1Hz with intensities ranging from 0.8-34 mW using a BioLED control module (model BLS-PL04-US) coupled with a High Power Fiber-Coupled LED Light Source (model BLS-FCS-0470-10) and Multimode Fiber Patchcords (Numerical aperture: 0.37 NA, Core size: 1000 μm. Catalog # FPC-1000-37-025MA, Mightex, Pleasanton, CA 94566, US). Following baseline optogenetic stimulation, isovalerate (200 μM) was applied to the mucosal epithelium for 15 min via a small metal ring placed over the receptive field of interest before afferent sensitivity to optogenetic stimulation was re-tested. Action potentials were analyzed offline using the Spike 2 wavemark function and discriminated as single units based on a distinguishable waveform, amplitude, and duration (CED, Cambridge, UK).

### Colon-pelvic nerve preparation and intact colon for whole nerve recordings

*To determine the effects of intraluminal isovalerate or CNO on* distension sensitive afferents, male and female wild type EC^hM3Dq^ or EC^PFTox^ mice. Preparations were similar to those described above for mucosal afferents; however, the preparation was kept intact, and the colon ligated at either end to allow for fluid distension (100 μL/min, 0-80mmHg). For wild type or EC^hM3Dq^ mice, distension was performed with CNO (100 μM) applied intraluminally. For wild type or EC^PFTox^ mice, distension was performed with isovalerate (200 μM) applied intraluminally. Whole pelvic nerve recordings were made using a sealed glass pipette containing a microelectrode (WPI, USA) attached to a Neurolog headstage (NL100AK; Digitimer, UK). Nerve activity was amplified (NL104), filtered (NL 125/126, bandpass 50–5,000 Hz, Neurolog; Digitimer, UK), and digitized (CED 1401; CED, Cambridge, UK) to a PC for offline analysis using Spike2 software (CED, Cambridge, UK). The number of action potentials crossing a pre-set threshold at twice the background electrical noise was determined per second to quantify the afferent response. Single-unit analysis was performed offline by matching individual spike waveforms through linear interpolation using Spike2 version 5.18 software. Afferent units sensitive to colonic distension were classified as Low Threshold, Wide Dynamic Range, or High Threshold based on their firing characteristics. Low threshold (LT) afferents responded below 10mmHg of distension with afferent firing rates at 20 mmHg >20% of the maximal response. Wide Dynamic Range (WDR) afferents responded below 10mmHg of distension with afferent firing rates at 20 mmHg <20% of the maximal response. High threshold (HT) afferents responded at distension pressures >10 mmHg.

### Intravital GCaMP imaging of L6 DRG

Male and female mice heterozygous for Na_V_1.8-Cre and GCaMP6s (Jax strain #024106; purchased from The Jackson Laboratory; Bar Harbor, ME, USA) were bred within SAHMRI’s specific and opportunistic pathogen-free animal care facility. Mice were then transported to the Core Animal Facility within the University of South Australia, where all procedures took place.

#### Surgical procedure

All surgical procedures, mice were anesthetized using 3% isoflurane in 100% oxygen at 2 L/min in an induction box before being transferred to a nose cone for maintenance at 1.5 – 2.5% isoflurane in 100% oxygen at 0.2 L/min. Anesthetized mice were kept on a heating pad to maintain body temperature. The surface above the region of interest was shaved and skin cleaned using alternating betadine and 70% ethanol. For the laminectomy procedure, an incision was made in the skin dorsal to the spinal region of interest, and the skin separated from the underlying muscle using spring-scissors. Using a scalpel, the fasciae of the exposed muscles were cut and retracted. Muscles were then separated from the vertebral column by blunt dissection using spring scissors, and any remaining muscle fragments were cleaned away. The dorsal laminectomy was performed by inserting the tips of small spring scissors between neighboring vertebrae, caudally to the vertebra of interest, and cutting. The transverse process was then removed, again using small spring scissors, to reveal the underlying DRG. Epaxial muscles from the left and right of the vertebral column were removed by blunt dissection to create space for the microscope objective to access the DRG. Any bleeding was controlled using gauze, cotton swabs, and warmed saline. Mice were placed in a spinal stabilization device to reduce movement artifacts due to breathing and heartbeat. Vertebrae rostral and caudal to the exposed DRG were clamped using two sets of forceps positioned ventrally to the transverse processes and forceps clamped within the device. Mice within the stabilization device were transferred to the confocal stage for imaging and maintained under isoflurane anesthesia.

#### Confocal imaging and intra-colonic administration

L6 DRGs were imaged on a Zeiss LSM710 confocal microscope equipped with both argon and helium-neon lasers. GCaMP6s fluorescence was captured using the argon laser, with excitation at 488 nm and collection at 493-556 nm. Images were captured using a Zeiss Plan-Apochromat 10x objective, 0.3 NA. Functional time-series were acquired at a rate of 0.985 frames per second, from an 850 × 850 μm region (512 × 512 pixels) using a pinhole aperture to image a slice thickness of 80 μm in the z-axis. Images were taken at baseline, following administration of 100 μl boluses of either intra-colonic i) saline, ii) isovalerate (200 μM), iii) alosetron (10 μM) or iv) isovalerate (200 μM) and alosetron (10 μM) combined.

#### Image processing

Drift and motion artifacts in time-lapse recordings were corrected using the motion correction module in the EZcalcium toolbox ^47^ implemented in MATLAB. Further image processing was completed using Fiji/ImageJ version 2.0.0 ^48^. Fluorescence traces of DRG cell bodies were extracted using custom software. Regions of interest (ROIs) were manually drawn around L6 DRG cell bodies, and their fluorescence traces were extracted as pixel averages. Baseline fluorescence (F0) was defined as the average fluorescence during the initial four frames, and traces were calculated as the change in fluorescence normalized to baseline (ΔF/F0). Drawing of ROIs and trace extraction was performed blind to the experimental group.

### Intestinal organoids

Adult EC^hM3Dq^, EC^PFTox^, and Tac^Cre^ male mice were used to generate intestinal organoids as previously reported ^49^. Organoids were maintained in organoid growth media (advanced Dulbecco’s modified Eagle’s medium/F12 supplemented with penicillin/ streptomycin, 10 mM HEPES, Glutamax, B27 [Thermo Fisher Scientific], 1 mM N-acetylcysteine [Sigma], 50 ng/mL murine recombinant epidermal growth factor [Thermo Fisher Scientific], R-spondin1 [10% final volume], and 100 ng/mL Noggin [Peprotech]) and were passaged every 6-7 days. For adeno-associated viral (AAV) infection of organoids, day 6-7 organoids were mechanically disrupted and re-plated into 50 μL of matrigel/AAV mixture (0.8 μL each of Cre-dependent AAV harboring tdTomato and GCaMP6s [Addgene, #28306-AAV9 and #100842-AAV9] mixed with 50 μL of matrigel [Corning]).

### ELISA

EC^hM3Dq^, EC^PFTox^, and Tac^Cre^ organoids were removed from matrigel, washed with cold DPBS + 0.1% BSA twice, and re-suspended into DPBS. Organoids were aliquoted into five tubes and incubated in 100 μL of Ringer’s solution (140 mM NaCl, 5 mM KCl, 2 mM CaCl2, 2 mM MgCl2, 10 mM D-glucose, and 10 mM HEPES-Na [pH 7.4]), Ringer’s + 20 μM allyl isothiocyanate (AITC), or Ringer’s + 1.7 μM deschloroclozapine (DCZ) for 15 min. The supernatant was collected and stored at −80°C. ELISA was performed according to the manufacturer’s protocol (LDN, #: BA E-5900R).

### Calcium imaging

AAV-infected organoids were mechanically dissociated 5-6 days post-infection and placed on Cell-Tak (Corning) coated coverslips in the imaging chamber containing Ringer’s solution. Organoids were maintained under a constant laminar flow of Ringer’s solution applied by a pressure-driven micro-perfusion system (SmartSquirt, Automate Scientific). EC cells were identified by tdTomato expression. GCaMP imaging was performed with an upright microscope equipped with an ORCA-ER camera and a Lambda LS (Sutter). GCaMP6s expression was confirmed before each measurement by applying high K+ solution (70 mM KCl, 70 mM NaCl, 2 mM MgCl2, 2 mM CaCl2, 10 mM D-glucose, and 10 mM HEPES-Na [pH 7.4]), and 1.7 μM DCZ was applied when the cell responded to high K+. ΔF/F0 was calculated after background subtraction.

### Behavioral experiments

All behavioral experiments were conducted between ZT6 – ZT10. All experimental mice were transferred to the behavior testing room 45 min prior to the beginning of the behavioral tests to habituate them to testing room conditions.

#### Elevated plus maze (EPM)

Mice were placed in the center of an elevated plus maze (EPM) 50 cm above the floor; the dimensions of each arm were 35 cm × 5 cm, with the closed arms having a 15 cm high wall and a center area of 5 cm × 5 cm. A mouse was placed in the center area of the maze and allowed to explore freely for 5 min. Each mouse was given one trial. The time spent and distance traveled in the open and closed arms were determined using video tracking software (ANY-maze, Stoelting, Wood Dale, IL, USA) and these measures served as an index of anxiety-like behavior. In DREADD experiments, DCZ injections (75 μg/kg) were administered intra-peritoneally for 10 mins before testing. For alosetron pre-treatment, mice were subcutaneously injected with 0.1 mg/kg 15 mins prior to DCZ injections.

#### Fear conditioning

Behavioral sessions were conducted in the Stoelting Fear Conditioning System consisting of fear conditioning chambers connected to tone and shock generators. Stimulus presentations were controlled by the ANY-maze software. On day 1 during fear acquisition, animals were acclimated to the conditioning chamber in context A for 3 minutes before exposure to four tone presentations that serve as the conditioned stimulus (CS, 75 dB for 30s). Each tone presentation co-terminated with an unconditioned stimulus (US, 0.5-mA foot shock for 2s). The inter-trial interval (ITI) was set at 2 mins. On day two during the context memory test, mice were exposed to context A for 6 minutes in the absence of any tone presentations. On day 3 during the fear memory test, mice were exposed to a new context (Context B) for 2 minutes followed by 4 CS presentations (with 1.5 min ITI). The % time spent freezing to context or tone was analyzed using ANY-maze software. Context A consisted of grid flooring and was cleaned with 0.02% sodium hypochlorite. Context B consisted of plexiglass flooring and was cleaned with 70% ethanol.

#### Marble burying

A standard 75 in^2^ cage containing clean wood chip bedding to a depth of 5 cm was prepared, with 15 marbles arranged uniformly on the surface. Mice were individually placed in the corner of the cage and the number of marbles buried (to 2/3 their depth) was counted after 30 min.

#### Nestlet shredding

Mice were placed in a standard cage with a pre-weighed nestlet. Nestlets were 5 mm thick cotton fiber weighing about ~3 g. Mice were left undisturbed for 90 min. The remaining unshredded nestlet was removed and weighed to determine the percentage of nestlet material shredded.

### Histology

Immunofluorescence (IF) in the small and large intestine was performed using 5 μm and 20 μm cryosections or whole-mount tissue as noted, in the brain and spinal cord using 25 um sections, and in DRGs using whole-mount tissue. Blocking was performed with 5% w/v BSA (Sigma), 5% normal serum corresponding to secondary antibody species in 0.3% Triton-X and PBS at room temperature for 60 minutes. Primary antibodies were incubated overnight at 4 °C at the indicated dilutions. Antibodies used were against serotonin (1:5000, Immunostar, 1:500 Abcam), GFP (1:500, Aves), and mCherry (1:500, Takara). Alexa Fluor-conjugated secondary antibodies were used at 1:500 (Millipore). Confocal imaging was performed on Nikon Ti2 microscope with Crest LFOV spinning disk and Nikon Ti microscope with Yokagawa CSU22 spinning disk. Images were assembled in ImageJ. Imaging was performed on the following systems: Nikon Ti2 microscope with Crest LFOV spinning disk, Nikon Ti microscope with Yokagawa CSU22 spinning disk, and BZ-X800 Keyence fluorescence microscope. Images were assembled in ImageJ.

### Gastrointestinal transit studies

Total gastrointestinal transit time was measured as previously described ^50,51^. Briefly, mice were fasted overnight on wire racks with free access to water. Carmine red 6% (w/v) solution in methyl cellulose was delivered by orogastric gavage (300 μL) and the time until the first red stool pellet recorded. For colonic transit, animals were fasted overnight on wire racks with free access to water. Animals were then lightly anesthetized with isoflurane and a 3mm diameter glass bead inserted in the colon to a depth of 2 cm. The time to bead expulsion was then recorded. For EC^hM3Dq^ experiments, DCZ (75 μg/kg i.p.) was administered to both Tac^Cre^ control and EC^hM3Dq^ expressing animals 15 min before the start of each study. Studies were standardized to begin ~9 A.M. local time.

### Serum serotonin measurements

Mice were anesthetized with isoflurane and whole blood was collected by submandibular venipuncture into an EDTA containing microtainer. Plasma was separated by centrifugation and snap frozen in liquid nitrogen. Samples were stored at −80 °C until analysis by the Vanderbilt Neurochemistry Core via Liquid Chromatography/Mass Spectrometry. For EC^hM3Dq^ experiments, a baseline sample was taken first and then one week later DCZ (75 μg/kg i.p.) was administered 15 min before the second blood draw. Blood draws were performed at 9 A.M. local time.

### Blinding and statistical analysis

All behavior, GCaMP, and nerve recordings were performed by blinded experimenters, with blinding codes resolved post-analysis to allow statistical comparisons. VMR data were normalized with the average maximal response taken as 100%. AUC data were analyzed using two-way ANOVA with Šidák’s multiple comparison test. The total AUC data were analyzed by unpaired two-tailed t-tests. Mucosal afferent data were analyzed to determine if they were distributed normally using Kolmogorov-Smirnov or Shapiro-Wilk tests, with subsequent analysis using two-way ANOVA with Šídák’s multiple comparisons tests or Wilcoxon matched-pairs signed rank tests. Optogenetic mucosal afferent data are expressed as the threshold for action potential activation (mW/ mm2) with significant differences determined by Chi-square tests. Colon-pelvic whole nerve data were analyzed using either two-way analysis of variance (ANOVA) or paired t-tests. Intravital GCaMP imaging data were analyzed using Student’s two-tailed unpaired t-tests. Gastrointestinal transit data were analyzed using Student’s t-test (unpaired, 2-tailed), whilst serum serotonin data were analyzed using Student’s t-test (unpaired, 2-tailed). Unless otherwise stated data are presented as mean ± SEM and analyzed using Prism 9 (GraphPad, San Diego, California USA, www.graphpad.com). Differences were considered statistically significant at P < 0.05. n represents the number of afferents/cells/ organoids and N represents the number of animals.

## ACKNOWLEDGMENTS

We thank Jeannie Poblete for expert assistance with animal husbandry and genotyping and Dr. N. Bellono for strategic contributions in the early stages of this project. We also thank the Garvan Institute, Australia for genotyping services and The University of South Australia and the UCSF Nikon Imaging Core for use of their in vivo confocal imaging facility. We thank Dr. K. Yackle and all members of our groups for many helpful suggestions and critical comments. This work was supported by NIH Training Grant T32 DK007762 and postdoctoral fellowship from the A.P. Giannini Foundation (R.D.M.), and the Damon Runyon Cancer Research Foundation (K.K.T.), a Simons Foundation Autism Research Initiative Pilot Award (514791 to D.J.), a Rainin Foundation Innovator Award (20191150 to D.J.), grants from the US National Institutes of Health (HEAL-SPARC initiative U01NS113869 to H.A.I., D.J. and S.M.B; R35 NS105038 to D.J.; R01 DK121657, GCRLE0320 to H.A.I., R03 DK121061, R01 DK128346 to J.R.B.), National Health and Medical Research Council of Australia (NHMRC) Investigator Leadership Grant (APP2008727 to S.M.B), an NHMRC Ideas Grant (APP1181448 to J.C.), and The Hospital Research Foundation (THRF) PhD Scholarship (SAPhD000242018 to J.M.).

## AUTHOR CONTRIBUTIONS

R.M. developed and implemented transgenic strategies for targeting DREADD and PFTox expression to EC cells. N.D.R., A.V. and R.D.M. characterized DREADD and PFTox expression in these lines histologically. J.C. and S.M.B. designed, performed, and analyzed ex vivo studies mechanically and optogenetically stimulating colonic mucosal afferents. M.B. and S.M.B. designed, performed, and analyzed ex vivo studies on colonic distension sensitive afferents. G.S., J.M., J.C., S.M.B, K.B., and J.R.B. designed, performed, and analyzed visceromotor response to colorectal distension. J.C., A.H. and S.M.B. designed, performed, and analyzed intravital GCaMP imaging studies. K.K.T. quantified serotonin release from organoids. A.V. performed and analyzed experiments to assess anxiety-related behaviors. C.B.S. collected blood samples for neurotransmitter analysis. H.A.I., J.R.B., S.M.B. and D.J. wrote the manuscript with input and suggestions from all authors and provided advice and guidance throughout.

## EXTENDED DATA INFORMATION (6 FIGURES)

**ED Fig1**: Isovalerate sensitizes mucosal afferents to optogenetic stimulation

**ED Fig 2**: Silencing EC cells attenuates responses to chemical irritants and noxious distension

**ED Fig 3:** Intersectional genetic strategy limits gene expression to gut EC cells

**ED Fig 4:** Activating EC cells increases afferent output and VMR to colorectal distension

**ED Fig 5**: Serum serotonin levels and gastrointestinal transit are unchanged in EC manipulation models

**ED Fig 6:** Activation and silencing of EC cells increases anxiety-like behaviors.

**ED Fig. 1.**
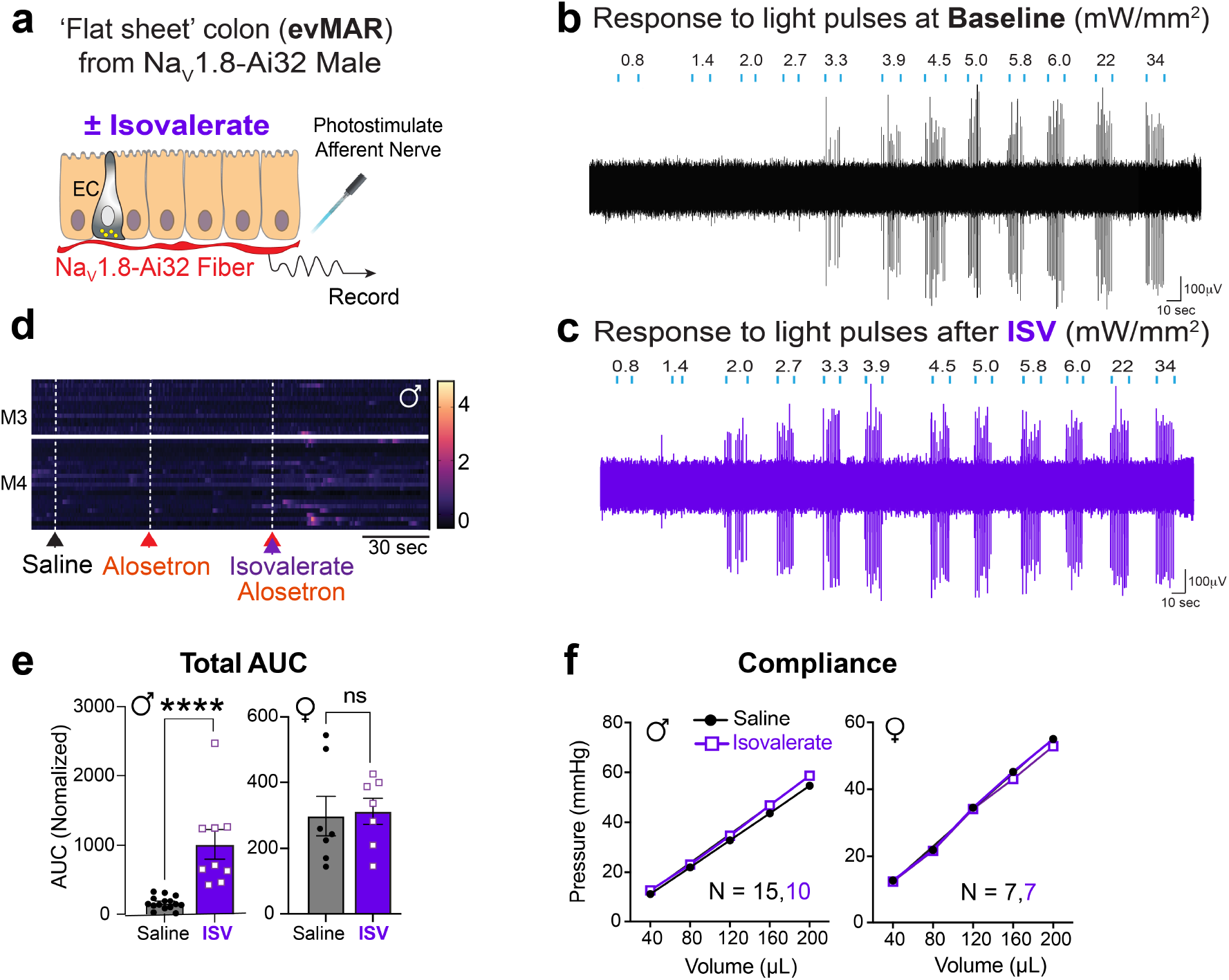
Sex-specific sensitization of mucosal afferents and visceral motor responses by isovalerate. **a,** Graphical scheme of ex vivo ‘flat sheet’ colon pelvic nerve *ex vivo* mucosal afferent recordings (evMAR) from Na_V_1.8-Ai32 mice. Mucosal afferents stimulated with light (l = 470 nm, 10 ms pulses at 1Hz with increasing intensity from 0.8 – 12 mW/mm^2^) plus vehicle or isovalerate (200 μM). **b-c,** Representative examples of pelvic mucosal afferent fiber recordings under (b) baseline and (c) isovalerate conditions in response to increasing intensities of light (0.8 – 34 mW/mm^2^) showing isovalerate lowers the threshold to optogenetic activation. **d,** Heatmaps from two representative males showing in vivo Na_V_1.8-GCaMP6s responses of L6 DRG neurons to intracolonic application of an isovalerate bolus (200 μM) after pretreatment with alosetron (10 μM). **e,** Group data showing total area under the curve for all colonic distension pressures showing VMRs are significantly increased in male but not female mice following intracolonic application of isovalerate (100 μl bolus of 200 μM). **f,** Colonic compliance is unchanged with isovalerate in both male and female mice. Wilcoxon matched-pairs signed-rank test in panel e and two-way ANOVA in panel f. ****p < 0.0001.

**ED Fig. 2.**
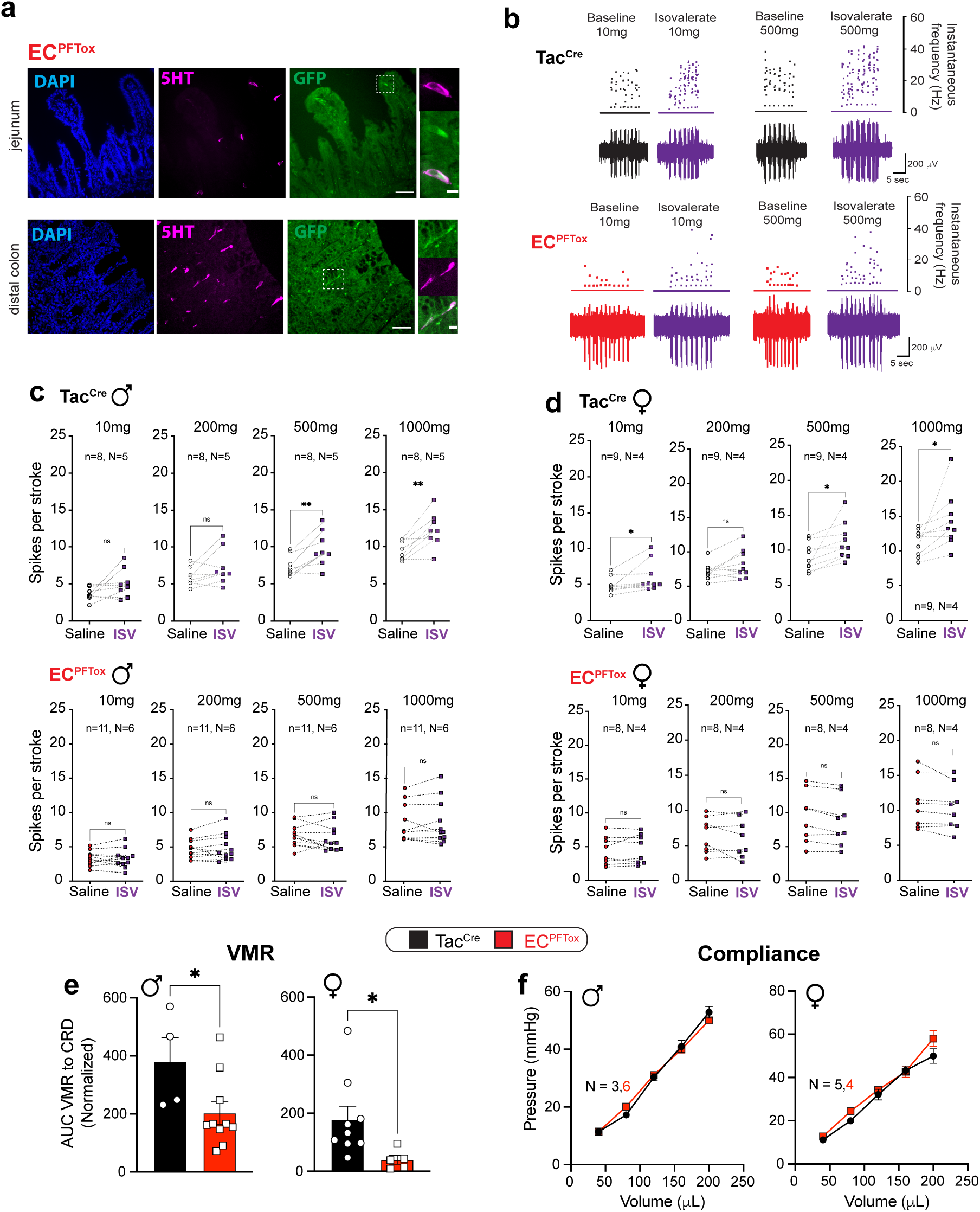
Silencing EC cells attenuates responses to chemical irritants and noxious distension. **a,** Histologic sections of EC^PFTox^ (*Pet1Flp;Tac1Cre;RC::PFTox*) in small and large intestine demonstrating colocalization of PFTox allele GFP reporter (green) and 5-HT (magenta). Scale bars 50 μm (left) and 10 μm (right). **b,** Representative examples of pelvic mucosal afferents firing more action potentials in response to stroking with 10mg or 500 mg von frey hairs (vfh) in the presence of isovalerate for control (upper) but not EC^PFTox^ (lower) mice. **c,d,** Group data showing before and after isovalerate (200 μM) application response to increasing mechanical stimulation with vfh for males (c) and females (d) for control (Tac^Cre^, upper panels) and EC^PFTox^ (lower panels) mice. **e,** Group data showing total area under the curve for all colonic distension pressures showing VMRs significantly reduced for both male (left) and female (right) EC^PFTox^ mice. **f,** Colonic compliance is unchanged in EC^PFTox^ animals. Wilcoxon matched-pairs signed-rank test and Student’s paired t-test in panels c, d. Student’s t-test (unpaired, 2-tailed) in panel e. *p <0.05.

**ED Fig. 3.**
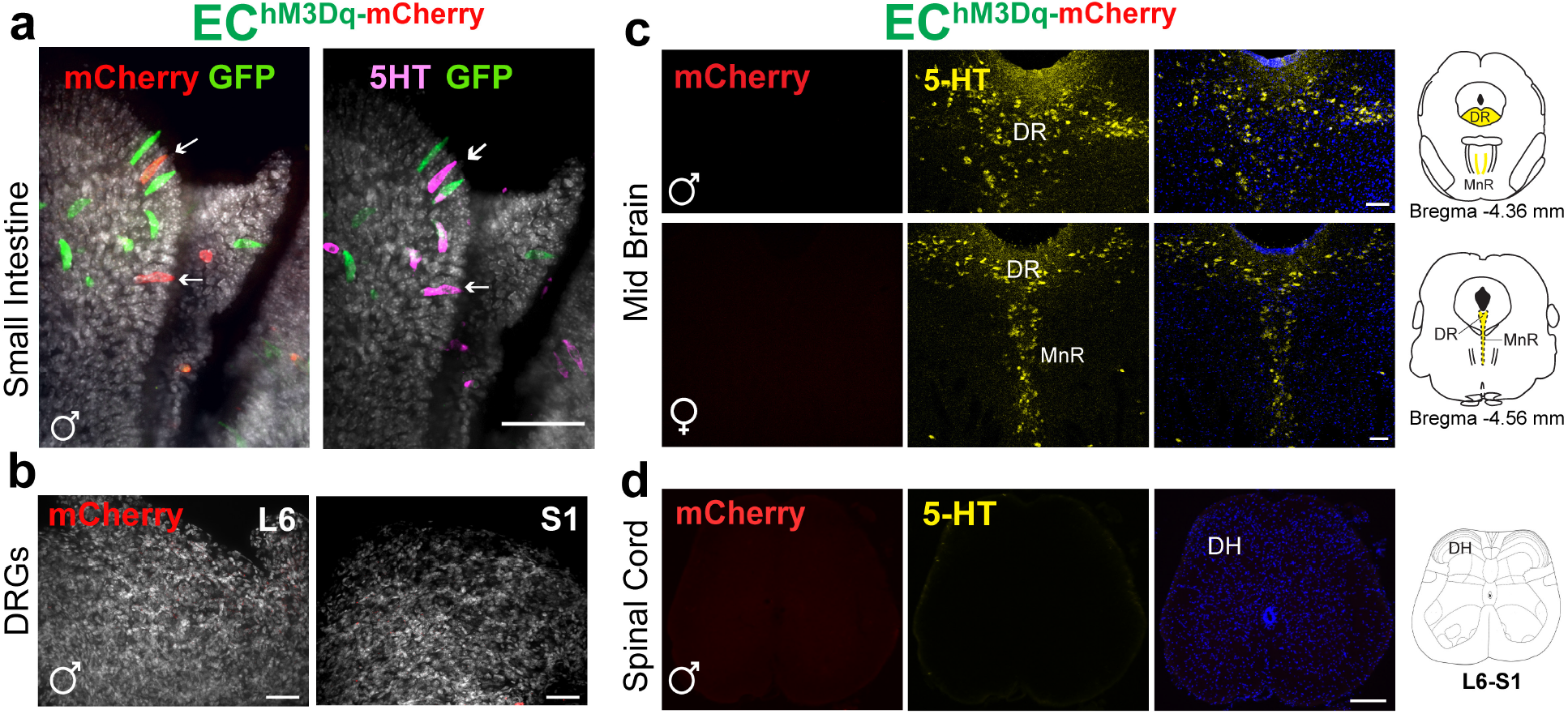
Intersectional genetic strategy limits gene expression to gut EC cells. **a,** Whole-mount small intestine (jejunum) of EC^hM3Dq^ (*Pet1FlpTac1Cre;RC::FL-hM3Dq*) demonstrating expression of single-(*Pet1Flp*, GFP, green) and double-recombination (*Pet1Flp/Tac1Cre*, mCherry, red) as indicated by hM3Dq allele reporters and 5HT (magenta). Scale bar 50 μm. **b,** Whole-mount DRG from spinal segments L6/S1 of EC^hM3Dq^ demonstrating the absence of mCherry (red) expression. Scale bars 50 μm. **c,** Representative images of EC^hM3Dq^ dorsal raphe (DR) and median raphe (MnR) nuclei showing 5HT-expressing neurons (yellow) and lack of mCherry (red) expression. **d,** Representative images of EC^hM3Dq^ lumbosacral spinal cord (L6-S1) showing absence of mCherry (red) and 5-HT (yellow) expression. Scale bar 100 μm, panel c and d.

**ED Fig. 4.**
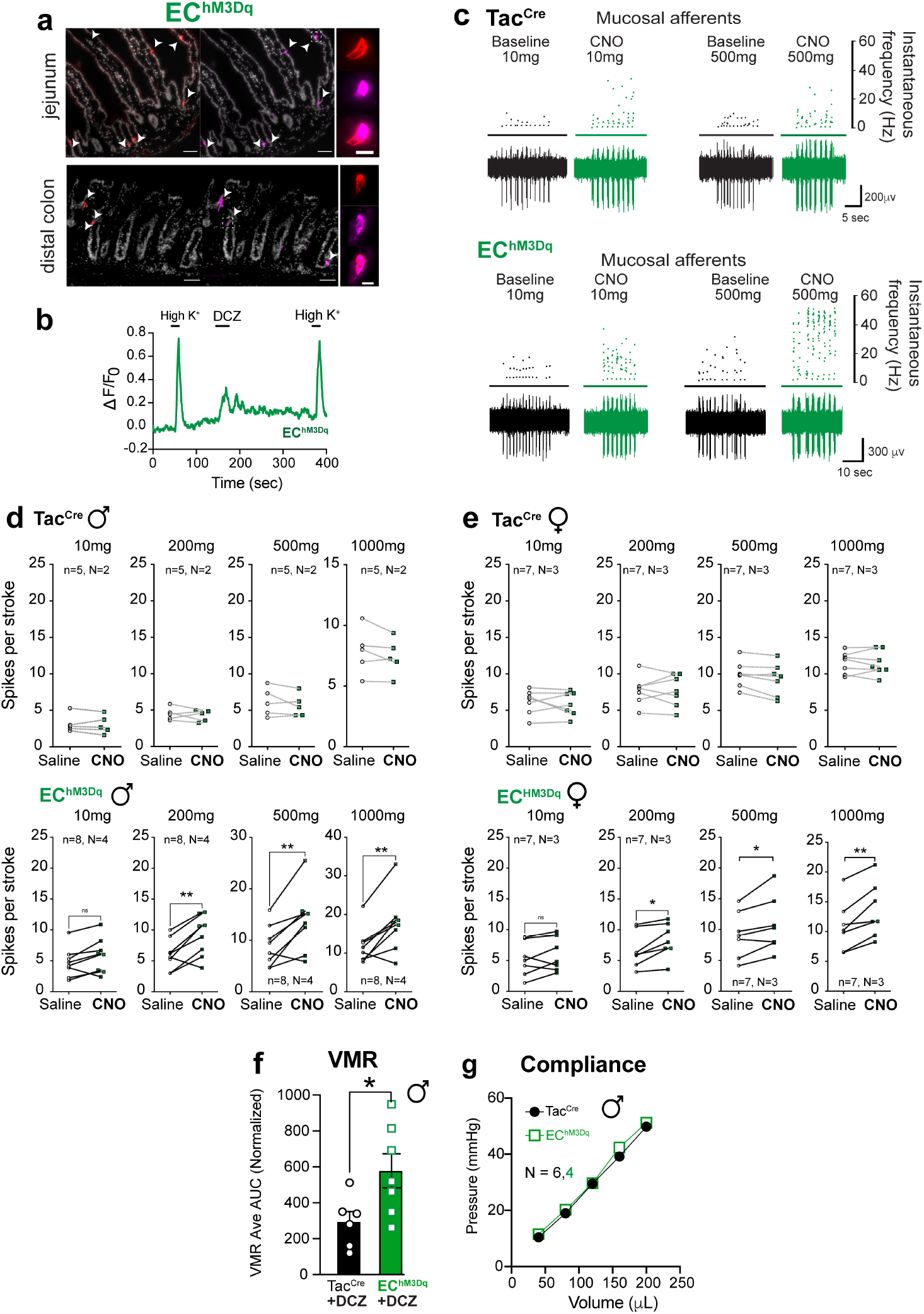
Activating EC cells increases afferent output and VMR to colorectal distension. **a,** Histologic sections from EC^hM3Dq^ (*Pet1Flp;Tac1Cre;RC::FL-hM3Dq*) mice showing small (jejunum) and large intestine demonstrating colocalization (white arrowheads) of the hM3Dq allele reporter (red) and 5-HT (magenta). Scale bars =50 μm (left) and = 10 μm (right). **b,** DREADD agonist DCZ (1.7 μM) elicits Ca2+ responses in EC^hM3Dq^ intestinal organoids as detected by change in fluorescence ratio. **c,** Representative examples of pelvic mucosal afferents firing action potentials in response to stroking with 10 mg, or 500 mg von frey hairs (vfh) in the presence of vehicle (black/grey) or CNO (100 μM; green) for control (left) and EC^hM3Dq^ (right) mice. **d, e,** Group data showing before and after CNO (100 μM) application response to increasing mechanical stimulation with vfh for males (d) and females (e) for control (Tac^Cre^ upper panels) and EC^hM3Dq^ (lower panels) mice. **f,** Group data showing total area under the curve for all colonic distension pressures showing VMRs significantly increased in male EC^hM3Dq^ mice following DCZ (75 μg/kg i.p.). **g,** Colonic compliance is unchanged in EC^hM3Dq^ animals. Wilcoxon matched-pairs signed-rank test in panels d and e. Student’s t-test (unpaired, 2-tailed) in panel f. *p <0.05.

**ED Fig. 5.**
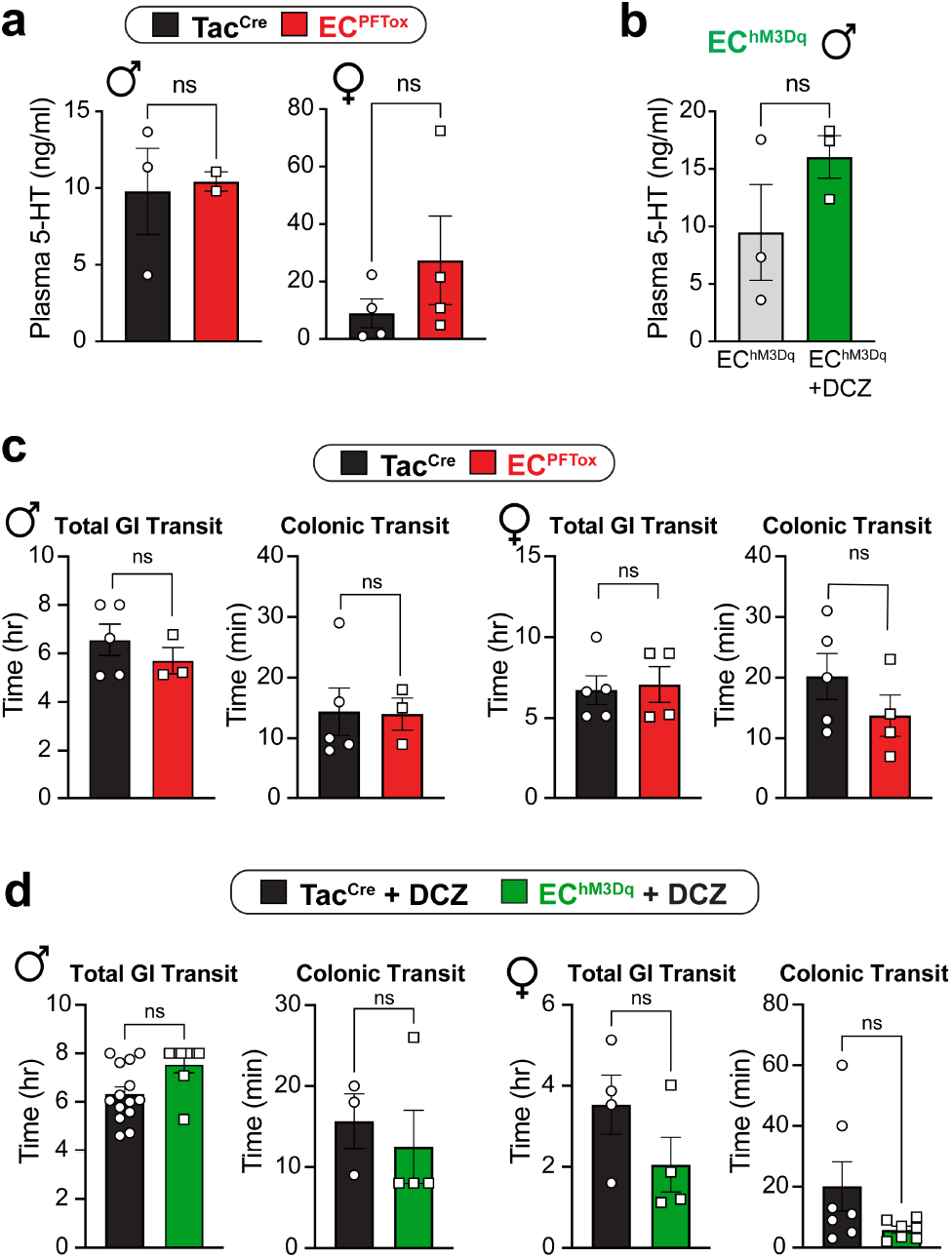
Serum serotonin levels and gastrointestinal transit are unchanged in EC manipulation models. **a,** Control (black) and EC^PFTox^ (red) serum 5-HT levels from male and female mice as determined by liquid chromatography/mass spectrometry (LC/MS). **b,** EC^hM3Dq^ Baseline (grey) and post-DCZ (75 μg/kg i.p.; green) 5-HT levels as determined by LC/MS. Post-DCZ serum was collected 15 min after i.p. injection. **c,** Total gastrointestinal and colonic transit times are similar between control (black) and EC^PFTox^ (red) mice. **d,** Total gastrointestinal and colonic transit times trend faster in DCZ-treated (75 μg/kg i.p.) EC^hM3Dq^ (green) mice relative to control (black) but are not statistically significant. Transit measurements started 15 min after i.p. injection. Statistics used Student’s t-test (unpaired, 2-tailed) for panels a-d.

**ED Fig. 6.**
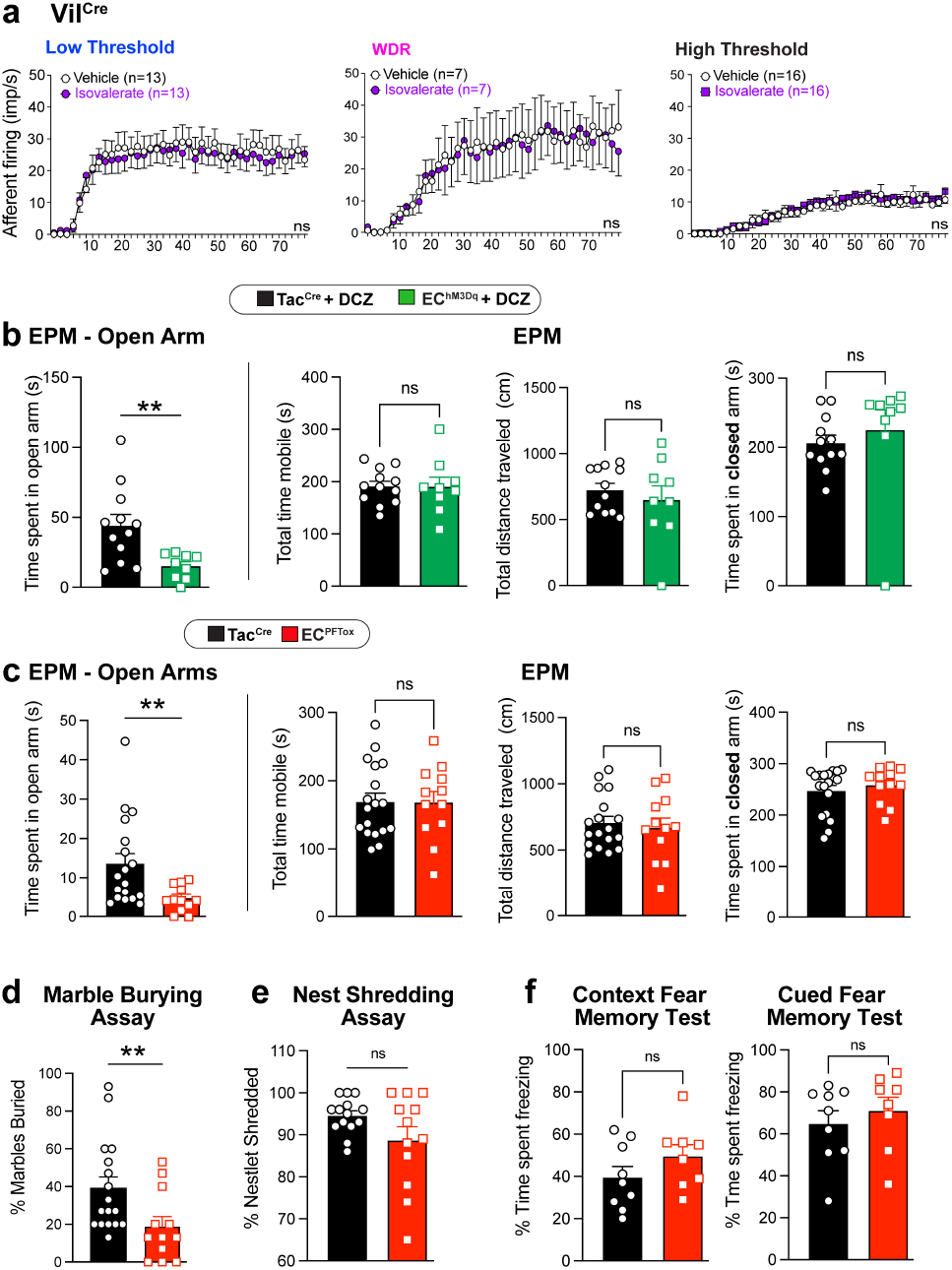
Activation and silencing of EC cells increase anxiety-like behaviors. **a,** Group data showing afferent firing to increasing distension pressures in colonic preparations from Tac^Cre^ mice at baseline and following isovalerate. **b,** Time spent in open arms of EPM following DCZ treatment (75 μg/kg i.p.) 10 min prior to testing. The total time mobile, total distance traveled, and total time spent in arms (both or closed) remain unchanged between EC^hM-3Dq^ and Tac^Cre^ control animals. **c,** Time spent in open arms of EPM for EC^PFTox^ animals. The total time mobile, total distance traveled, and total time spent in arms (both or closed) remain unchanged between EC^PFTox^ and Tac^Cre^ animals. **d,** EC^PFTox^ mice show significantly reduced marble-burying behavior compared to Tac^Cre^ controls. **e,** EC^PFTox^ mice do not show differences in nestlet shredding behavior. **f,** EC^PFTox^ mice do not show deficits in contextual and cued fear conditioning. Two-way ANOVA (Šidák’s multiple-comparisons test) for panel a. Student t-test (unpaired Mann Whitney U test, 2-tailed) for panels b-f. **p < 0.01.

